# *Ex vivo* delivery of regulatory T cells for control of alloimmune priming in the donor lung

**DOI:** 10.1101/2021.02.07.430098

**Authors:** Ei Miyamoto, Akihiro Takahagi, Akihiro Ohsumi, Tereza Martinu, David Hwang, Kristen M. Boonstra, Betty Joe, Juan Mauricio Umana, Ke F. Bei, Daniel Vosoughi, Mingyao Liu, Marcelo Cypel, Shaf Keshavjee, Stephen C. Juvet

**Affiliations:** Latner Thoracic Surgery Research Laboratories, University Health Network, University of Toronto, Toronto, Ontario, Canada

## Abstract

Survival after lung transplantation (LTx) is hampered by uncontrolled inflammation and alloimmunity. Regulatory T cells (Tregs) are being studied for post-implantation cell therapy in solid organ transplantation. Whether these systemically administered Tregs can function at the appropriate location and time is an important concern. We hypothesized that *in vitro* expanded, recipient-derived Tregs can be delivered to donor lungs prior to LTx via *ex vivo* lung perfusion (EVLP), maintaining their immunomodulatory ability.

In a rat model, Wistar Kyoto (WKy) CD4^+^CD25^high^ Tregs were expanded *in vitro* prior to EVLP. Expanded Tregs were administered to Fisher 344 (F344) donor lungs during EVLP; left lungs were transplanted into WKy recipients. Treg localization and function post-transplant were assessed. In a proof-of-concept experiment, cryopreserved expanded human CD4^+^CD25^+^CD127^low^ Tregs were thawed and injected into discarded human lungs during EVLP. Rat Tregs entered the lung parenchyma and retained suppressive function. Expanded Tregs had no adverse effect on donor lung physiology during EVLP; lung water as measured by wet-to-dry weight ratio was reduced by Treg therapy. The administered cells remained in the graft at 3 days post-transplant where they reduced activation of intragraft effector CD4^+^ T cells; these effects were diminished by day 7. Human Tregs entered the lung parenchyma during EVLP where they expressed key immunoregulatory molecules (CTLA4^+^, 4-1BB^+^, CD39^+^, and CD15s^+^). Pre-transplant Treg administration can inhibit alloimmunity within the lung allograft at early time points post- transplant. Our organ-directed approach has potential for clinical translation.

## Introduction

Survival after lung transplantation (LTx) is limited by chronic lung allograft dysfunction, which is driven primarily by the recipient ‘s alloimmune response. CD4^+^CD25^hi^Foxp3^+^ regulatory T cells (Tregs) can inhibit allograft rejection by various mechanisms, and are currently being studied as a post-implantation adoptive cellular therapy in clinical trials of kidney and liver transplantation [1]. Although secondary lymphoid organs are an important site of immune regulation by Tregs, it has been known for many years that the presence of a high Treg-to-conventional T cell (Tconv) ratio within allografts is required for allograft acceptance [2, 3]. Moreover, animal data indicate that lung allografts can be rejected in the absence of secondary lymphoid organs – and that the initial priming of alloreactive T cells occurs within the lung allograft itself [4, 5]. These observations raise the concern that post-transplant systemic Treg administration in LTx recipients may not provide timely immune regulation at relevant sites *in vivo*.

We have developed and translated to the bedside, a technique of extended *ex vivo* lung perfusion (EVLP) as a platform for donor lung assessment and treatment [6]. We have also demonstrated successful to delivery of gene and cellular therapy in animal and human lungs [7, 8]. Pre-transplant administration of recipient-derived Tregs directly into donor lungs affords the unique opportunity to seed the allograft with therapeutic immune regulatory cells in advance of the arrival of alloreactive T cells after graft implantation. Since Tregs can be expanded, cryopreserved and thawed at a later date while retaining function [9], this approach is suitable for LTx from deceased donors.

Here, we tested the hypothesis that *in vitro* expanded recipient Tregs administered to the donor lung allograft during EVLP prior to transplantation creates an immunoregulatory environment in the donor lung leading to a diminished immune response within the lung allograft post-implantation. We used a rat EVLP-to-LTx experimental model to examine how Tregs interact with the lungs and how they may inhibit immune responses in the recipients. In a proof-of- concept experiment, we showed that cryopreserved expanded human Tregs can be delivered successfully to human lungs on EVLP.

## Materials and Methods

### Expanded rat Treg injection during rat EVLP followed by LTx

To study EVLP-based Treg therapy, we opted for a rat model amenable to EVLP followed by single LTx, based on our and others ‘ experience [10, 11]. We used a Fisher 344 (F344, RT1^lv^)- to-Wistar Kyoto (WKy, RT1^l^) strain combination [12]. Our protocol is depicted in figure 1a. WKy CD4^+^CD25^high^ cells were isolated by magnetic and fluorescence-activated cell sorting and expanded with anti-CD3 and anti-CD28 coated beads (Miltenyi Biotec) and 1000 units/mL recombinant human IL-2 (rhIL-2, Chiron) for 7 days. Expanded Tregs were labelled with CMTMR and/or eF450 cell tracker dye and resuspended in 1 mL perfusate solution (Steen®, XVIVO Perfusion, Denver, CO). Normothermic acellular EVLP was performed as previously reported [10]. A total of 4.3∼211.3 × 10^6^ live Tregs/kg F344 donor body weight were injected into the EVLP circuit upstream of the lungs at 60 min of EVLP. In control experiments, grafts received 1 mL Steen containing no cells. Perfusate was sampled upstream and downstream of the lung. At 180 min after injection, the right lung was used for end-EVLP analysis and the left lung was transplanted into a WKy recipient. Recipients were euthanized at days 3 or 7 post-transplant. The animal study was performed in accordance with the policies formulated by the Canadian Council on Animal Care and the protocol was approved by the Institutional Animal Care Committee (AUP2853).

**Figure 1.**
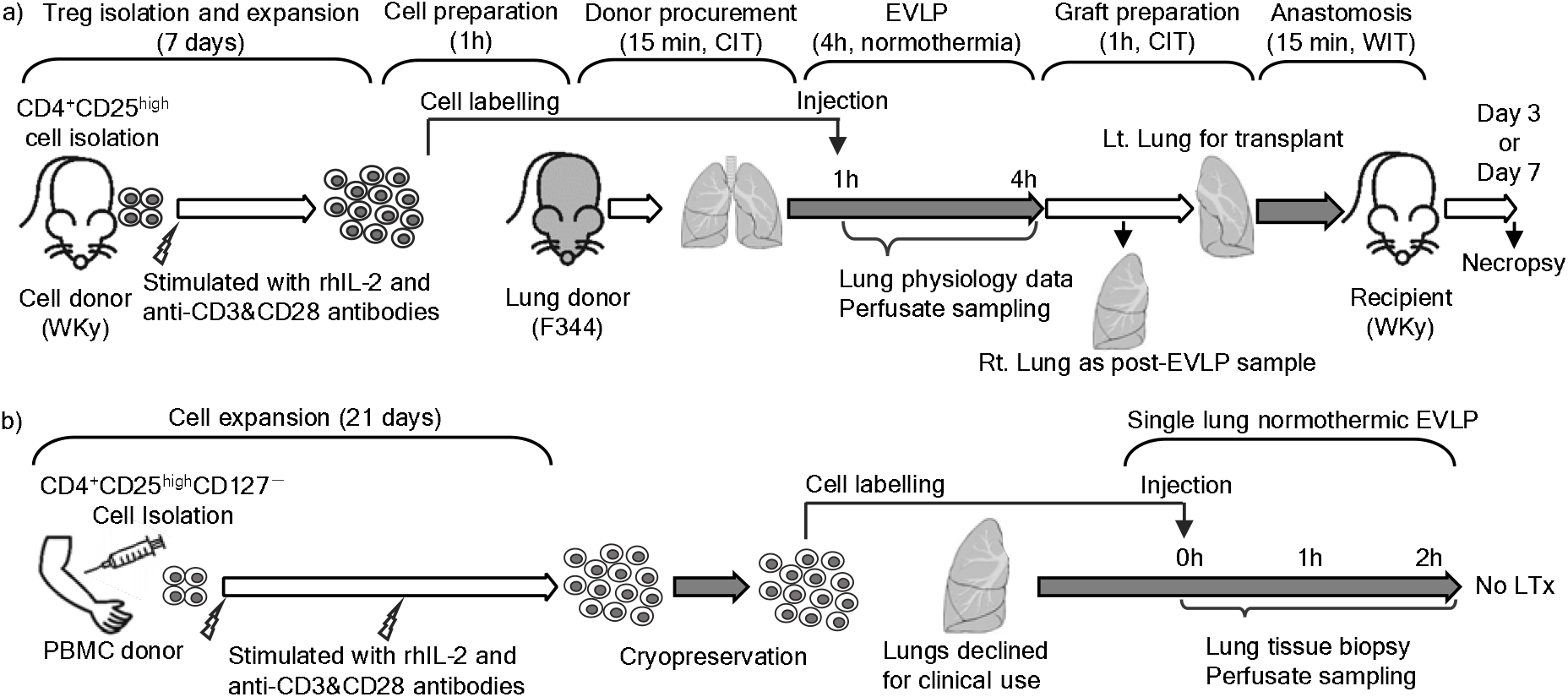
Study protocols. Schematic of rat Treg experiment. CD4^+^CD25^high^ cells (supplementary figure 1a) were isolated from WKy rat lymph nodes and expanded with rhIL-2 and anti-CD3 and CD28 antibodies for 7 days. Prior to starting EVLP with a F344 rat heart and lung block, the expanded cells were dye labelled for injection. Labelled Tregs or perfusate vehicle were injected to the circuit at the PA port 1 hour after the start of EVLP. After 4 hours, the left lung graft was transplanted to a WKy recipient while the right lung was subjected to further analyses. (b) Schematic of human Treg experiment. CD4^+^CD25^high^CD127^low^ cells from a healthy blood donor were isolated and expanded *in vitro*. After 21 days, the expanded Tregs were cryopreserved in liquid nitrogen. Upon notification that a lung was declined for transplantation on EVLP, Tregs were rapidly thawed, labeled with CMTMR and eF450, washed and injected into the EVLP circuit (supplementary figure 6b). EVLP continued for up to 2 hours after Treg injection, at which point tissue and perfusate were analyzed. Treg, regulatory T cell; EVLP, *ex vivo* lung perfusion; WKy, Wistar Kyoto; F344, Fisher 344; rhIL-2, recombinant human IL-2; PBMC, peripheral blood mononuclear cells.

### Expanded human Treg injection during human EVLP

The protocol for the human experiment is shown in figure 1b. CD4^+^CD25^+^CD127^low^ cells were isolated from a healthy donor and expanded for 21 days using anti-CD3 and anti-CD28-coated beads (Miltenyi Biotec) and 300 units/mL rhIL-2 (Chiron). At day 21, Tregs were cryopreserved. Human donor lungs on normothermic acellular EVLP deemed unsuitable for transplantation due to alveolar edema and/or poor compliance were used for the study (n = 3). Left or right single lung EVLP was established by truncating or clamping the hilum of the worse lung. Cryopreserved human Tregs were thawed and labelled with CMTMR and eF450 cell tracker dye. A total of 0.4∼0.8 × 10^9^ Tregs were resuspended in 20 mL Steen and injected into the EVLP circuit upstream of the lung. Perfusate and lung tissue were sampled just before Treg injection and at 60 min and 120 min after injection. Tissue samples taken at similar time points from contemporaneous declined lungs undergoing EVLP (no Treg injection, n = 5) were also obtained as controls. Experiments on human lungs were performed in accordance with the Helsinki declaration and was approved by the Institutional Research Ethics Board (06-283).

## Results

### Characteristics of isolated and expanded rat Tregs

We sorted WKy CD4^+^CD25^high^ cells (highest 2% of CD25^+^ CD4^+^ T cells, supplementary figure 1a) and found that they were 73.4 ± 12.8% Foxp3^+^. Expansion with anti-CD3 and anti-CD28- coated beads and IL-2 resulted in 152.8 ± 17.1-fold expansion (figure 2a, n = 21), with largely retained Foxp3 expression (63.5 ± 8.3% FoxP3^+^, supplementary figure 1b) and CD25 expression (93.6 ± 1.5% CD25^+^) at day 7. They also upregulated CD8 (81.3 ± 2.4% CD8^+^) at day 7, as reported previously [13]. The expanded cells dose-dependently suppressed *in vitro* proliferation of polyclonally stimulated CD4^+^CD25^low^ Tconv (figure 2b).

**Figure 2.**
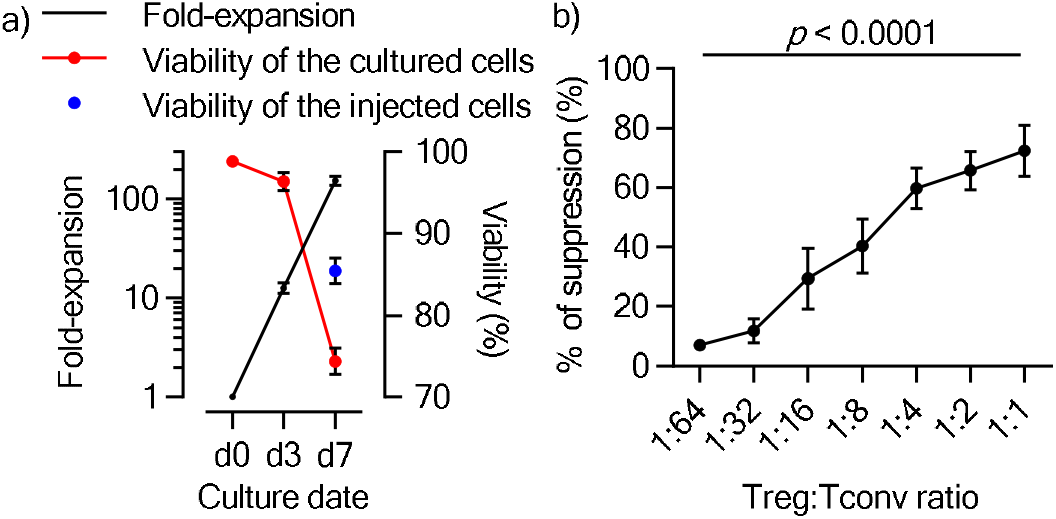
*In vitro* expansion of Tregs with keeping immunosuppressive effect. (a) Treg expansion. Fold-increase and viability of expanded Tregs on day 3 and day 7 of culture shown. Expanded cells were washed, increasing the viable proportion of injected cells to > 80%. (b) WKy Tregs mediate dose-dependent suppression of autologous Tconv (*p* < 0.0001, ANOVA). Gating strategy to identify CFSE-labelled Tconv is shown in supplementary figure 1c. Treg, regulatory T cell; Tconv, conventional T cells.

### Interaction of Tregs with lung allografts during rat EVLP

Tregs were tracked during EVLP (supplementary figure 1d-e) by dye labeling (figure 3a). The proportion of Tregs among live cells in both pulmonary artery (PA) and vein (PV) perfusate rapidly increased after injection and plateaued within 60 min (figure 3b). Remarkably, roughly 25% of administered Tregs remained in the EVLP circuit at the end of EVLP regardless of cell dose (figure 3c). Treg infusion did not affect lung compliance (figure 3d), vascular resistance (figure 3e), or mean PA pressure (figure 3f). Gas exchange – as measured by the ratio of the arterial partial pressure of oxygen to the fraction of inspired oxygen (P/F ratio, figure 3g), and perfusate glucose and lactate concentrations (supplementary figure 2a-b) were unaffected by Treg infusion.

**Figure 3.**
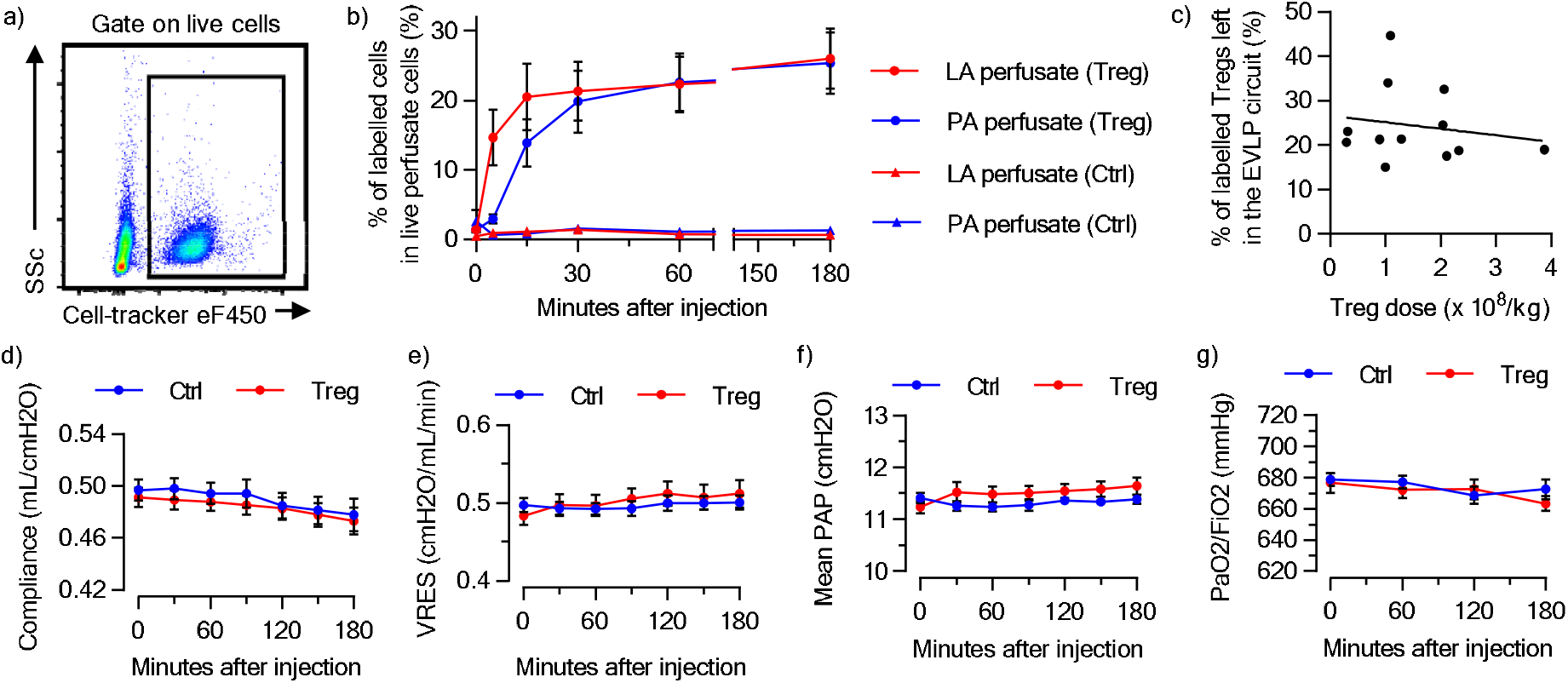
Administration of expanded WKy Tregs to allogeneic F344 lungs during EVLP. (a) Identification of labelled Tregs in EVLP perfusate by flow cytometry. (b) Proportion of Tregs in live perfusate cells at the PA and LA ports over time, which equalized at 60 minutes after injection. (c) The percentage of Tregs remaining in the circuit at the end of EVLP was calculated as 100 × (live cell count in the perfusate) × (fraction of Tregs in live perfusate cells)/(number of injected cells). (d) Compliance, (e) Vascular resistance (VRES), (f) Mean pulmonary arterial pressure (PAP) of the donor lungs on EVLP, and (g) ratio of the partial pressure of oxygen in the perfusate (PaO2) to the fraction of inspired oxygen (FiO2) of allogeneic F344 lungs during EVLP with or without Treg injection. WKy, Wistar Kyoto; Treg, regulatory T cell; F344, Fisher 344; EVLP, *ex vivo* lung perfusion; PA, pulmonary artery; LA, left atrium.

### Administered Tregs enter lung allografts and reduce lung vascular permeability at end of EVLP

There was no difference in acute lung injury scores at the end of EVLP between Treg-treated lungs and controls (figure 4a-b). The wet-to-dry weight ratio of the lung graft at the end of EVLP was lower in Treg-treated lungs (figure 4c), suggesting that Tregs may have helped to maintain alveolar integrity; despite this observation, ZO-1 expression was similar among Treg-treated lungs and controls (supplementary figure 2d).

**Figure 4.**
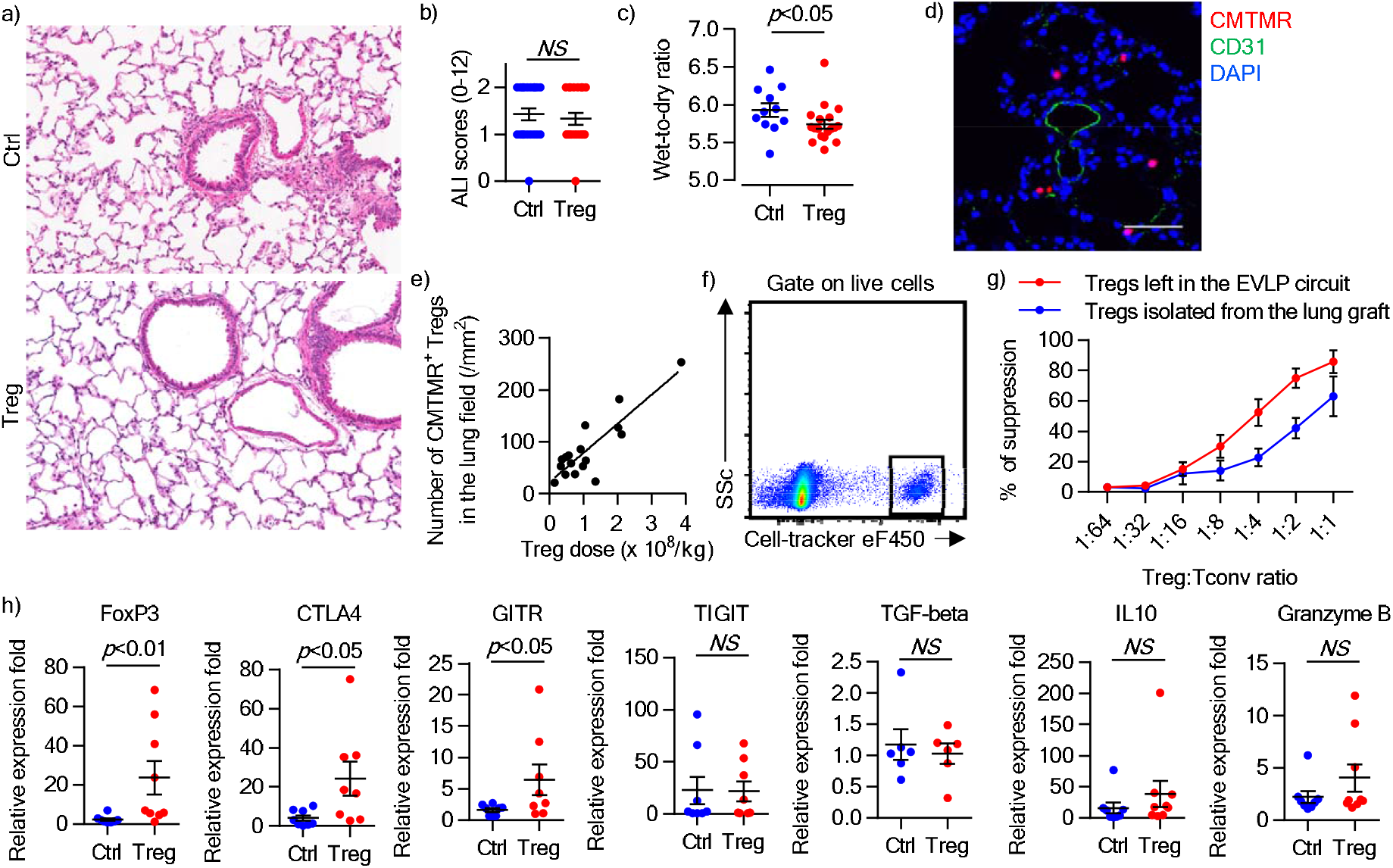
Expanded WKy Tregs enter allogeneic F344 lungs during EVLP while maintaining their regulatory function and phenotypic properties. (a) Representative histology of the lung graft after EVLP (hematoxylin and eosin staining; × 40). (b) Acute lung injury (ALI) score. Semi-quantification of alveolar space hemorrhage, vascular congestion, interstitial edema/fibrin deposition, and interstitial infiltration of white blood cells is shown in supplementary figure 2c. (c) Wet-to-dry weight ratio in Treg-treated and control lungs. Control grafts were significantly heavier than Treg-treated ones (*p* < 0.05, Mann-Whitney U test). (d) Immunofluorescence staining of the lung graft after EVLP demonstrates CMTMR^+^ (red) Tregs located in the lung tissue outside CD31^+^ (green) vessels. Scale bar=25 µm. (e) The number of CMTMR^+^ cells in the lung parenchyma was correlated with the injected dose of Tregs (*p* < 0.0001, R^2^ = 0.7542, Y = 0.5580 × X + 22.75). (f) Identification of transferred Tregs in the digested lung by flow cytometry. (g) Tregs sorted from perfusate and digested lung tissue at the end of EVLP exhibited dose-dependent suppressive activity toward autologous Tconv (*p* = 0.0001 and *p* < 0.0001 for Tregs isolated from the lung graft and those left in the EVLP circuit, respectively; one-way ANOVA). (h) Treg-related transcripts in the lung graft after EVLP, represented as a relative expression fold based on house-keeping gene. FoxP3, CTLA4, and GITR expression were significantly increased in the lung graft of Treg-treated animals compared to control cases. WKy, Wistar Kyoto; Treg, regulatory T cell; F344, Fisher 344; EVLP, *ex vivo* lung perfusion; Tconv, conventional T cells.

At the end of EVLP, CMTMR^+^ Tregs could be seen outside CD31^+^ vascular structures in the lung parenchyma (figure 4d). Tregs entered the lung in a dose-dependent manner, without evidence of saturation in the range of doses tested (figure 4e). We sorted CMTMR^+^ eF450^+^ Tregs from digested lung tissue at the end of EVLP (figure 4f) and found that they could suppress the proliferation of syngeneic polyclonally-stimulated Tconv to an extent that was comparable to, albeit slightly lower than, that of Tregs remaining in the EVLP circuit (figure 4g). Compared to control lungs, Treg-treated lungs, as expected, had higher levels of the Treg-related transcripts FoxP3, CTLA4, GITR, and CCR4 at the end of EVLP (figure 4h). There was no difference in the rate of apoptosis, as measured by TUNEL staining, in Treg-treated lungs up to 200 million Tregs per kg donor body weight, compared to controls (supplementary figure 2e). Further, Tregs themselves were not apoptotic at the end of EVLP (0.97 ± 0.52%, supplementary figure 2f).

### Transferred Tregs remain functional in the recipient post-transplant

Transferred Tregs were still detectable in the lung graft on day 3 post-transplant (figure 5a). Compared to the end of EVLP, the transferred cells had shifted to a predominantly subpleural, rather than a perihilar location (figure 5b). Tregs mediate contact-dependent modulation of MHC class II^+^ antigen-presenting cells [14]. Interestingly, the percentage of Tregs adjacent to MHC class II-expressing cells in the lung graft was increased on day 3 compared to at the end of EVLP (figure 5c). Further, FoxP3 expression by CD4^+^ T cells was higher in Treg-treated allografts than in controls at day 3 post-transplant (figure 5d). Transferred Tregs were also identified in the draining lymph nodes (dLNs) on day 3, within the extravascular space (figure 5e).

**Figure 5.**
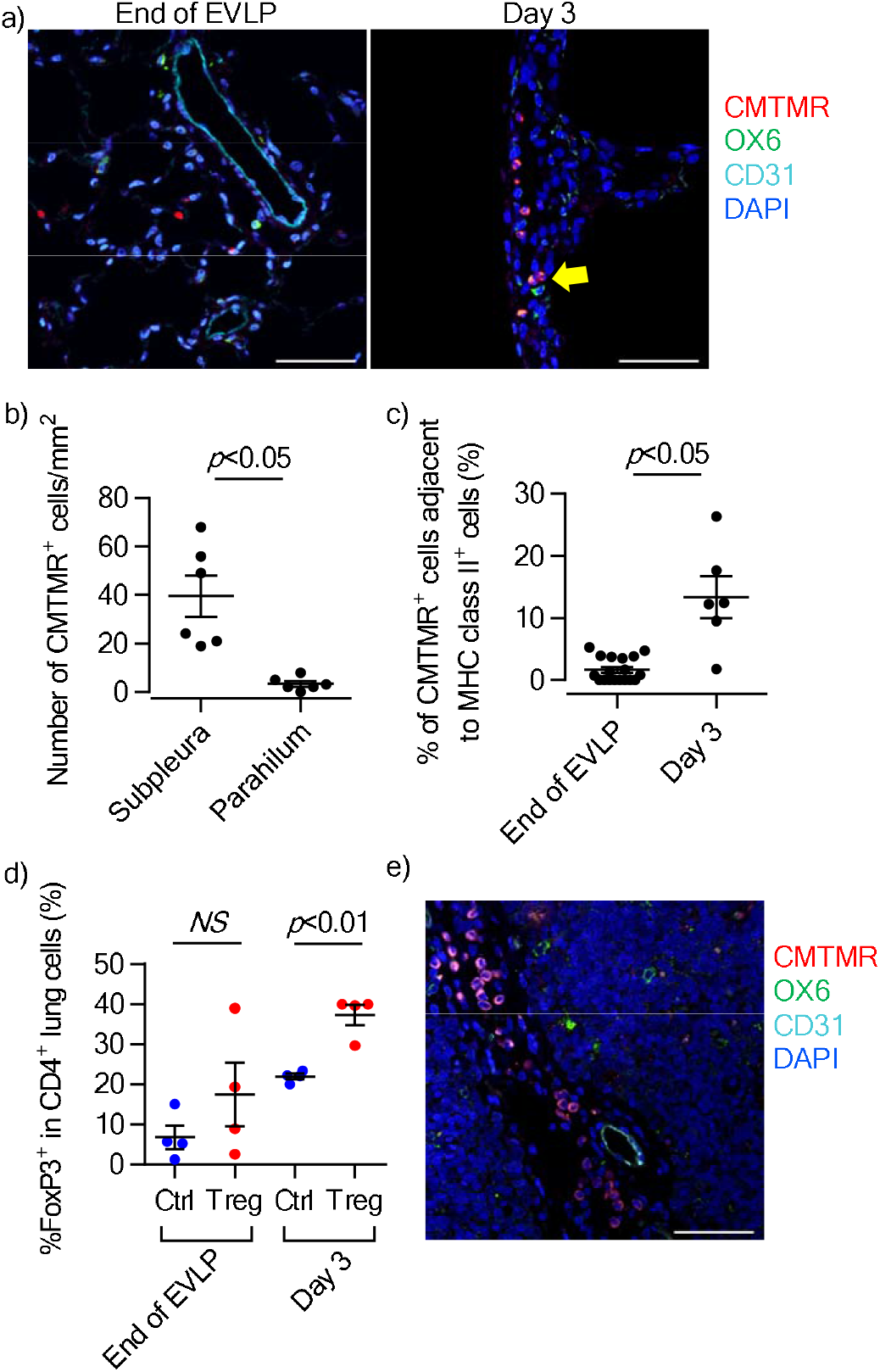
Tracking Tregs in the recipient after lung transplantation. (a) Compared to at the end of EVLP, on day 3 post-transplant transferred CMTMR^+^ Tregs (red) were predominantly located in the subpleural area (< 50 µm from pleural surface) and often seen in proximity to MHC class II^+^ (green) cells (yellow arrow). Images from each channel are shown in supplementary figure 3a-b. Scale bars = 50 µm. (b) Spatial distribution of Tregs in the graft on day 3 (*p* < 0.05, Mann-Whitney U test), and (c) increased proximity of Tregs to MHC class II^+^ cells on day 3 (*p* < 0.05, Mann-Whitney U test). (d) The proportion of intragraft CD4^+^ T cells expressing Foxp3 was increased in Treg-treated grafts compared to controls at day 3 and day 7 post-transplant (*p* < 0.01, Welch ‘s T test). Gating strategy to identify intragraft CD4^+^ T cells and the representative histograms of FoxP3 expression in them are shown in supplementary figure 3c and supplementary figure 3d, respectively. (e) Identification of CMTMR^+^ Tregs (red) in the mediastinal lymph nodes of recipients of Treg-treated lung allografts on day 3. Images from each channel are shown in supplementary figure 3e. Scale bar = 50 µm. Treg, regulatory T cell; EVLP, *ex vivo* lung perfusion; MHC, major histocompatibility complex.

At day 3 post-transplant, there was no meaningful difference in acute lung injury scores between Treg-treated animals and controls, with allografts in both groups exhibiting only mild abnormalities (figure 6a-b). The number of CD3^+^ Tconv cells adjacent to MHC class II^+^ cells was reduced in Treg-treated lung grafts compared to controls on day 3 (figure 6c-d). Moreover, within Treg-treated lung allografts, fewer CD3^+^ Tconv cells were adjacent to MHC class II^+^ cells that were closely associated with transferred Tregs in comparison with those that were not (figure 6e).

**Figure 6.**
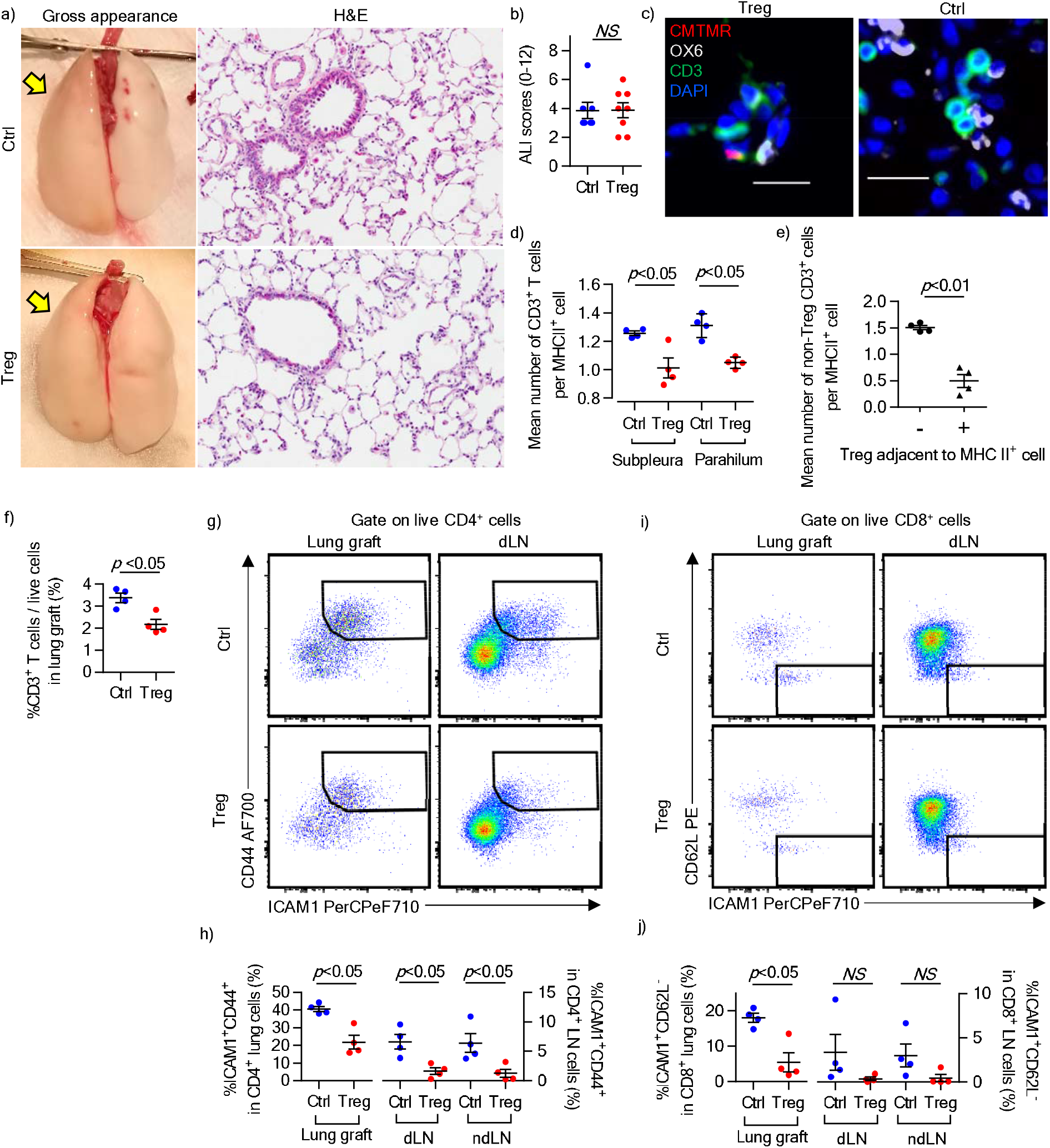
Post-transplant immune regulation by expanded Tregs delivered to the allograft prior to transplantation. (a) Representative pictures of the gross appearance and the histology (hematoxylin and eosin staining; × 40) of the lung graft (yellow arrows) on day 3 post-transplant. (b) Acute lung injury (ALI) scores in Treg-treated and control grafts. Details of histological semi-quantification are shown in supplementary figure 4a. (c) Representative high power immunofluorescence images showing intra-graft MHC class II^+^ cells (white) adjacent to CMTMR^-^CD3^+^ non-transferred Tconv (green) and CMTMR^+^CD3^+^ transferred Tregs (red). Scale bars=20 µm. (d) Mean number of CD3^+^ T cells adjacent to MHC class II^+^ cells in at least 10 high powered fields in Treg-treated and control lungs at the subpleural and perihilar areas. (e) Number of Tconv adjacent to MHC class II+ cells without (-) or with (+) an adjacent Treg cell (*p* < 0.01, Mann-Whitney U test). (f) Percentage of CD3^+^ T cells (gating strategy shown in supplementary figure 4b), as well as CD4^+^ T cells and CD8^+^ T cells (supplementary figure 4c), was significantly decreased in Treg-treated lung graft on day 3. (g) Flow cytometric analysis of CD44 and ICAM1 expression on CD4^+^ cells from the lung allograft and dLN. (h) Percentage of ICAM1^+^CD44^+^ cells in the CD4^+^ T cell compartment in the lung, dLN and ndLN at day 3. (i) Flow cytometric analysis of CD62L and ICAM1 expression on CD8^+^ cells from the lung allograft and dLN. (j) Percentage of ICAM1^+^CD62L^-^ cells in the CD8^+^ T cell compartment in the lung, dLN and ndLN at day 3. Treg, regulatory T cell; MHC, major histocompatibility complex; Tconv, conventional T cells; dLN, draining lymph node; ndLN, non-draining lymph node.

The percentage of CD3^+^ T cells among live cells at day 3 post-transplant was decreased by Treg treatment (figure 6f). Upregulation of ICAM1 and CD44 and downregulation of CD62L have been reported as rat Tconv activation markers [15–17]. We observed activated Tconv in the lungs and dLNs at day 3 (supplementary figure 4d-i). Treg administration during EVLP reduced the population of activated CD4^+^ T cells expressing ICAM1 and CD44 in both the graft and dLNs. Examination of other activated subsets, namely the ICAM1^+^CD62L^-^ population (supplementary figure 4j-k) and the CD44^+^CD62L^-^ population (supplementary figure 4l-m) revealed similar findings. The population of activated ICAM1^+^CD62L^-^ CD8^+^ T cells was also decreased by Treg in the treated lung graft, but not in the dLNs (figure 6i-j). The expression of FoxP3 and Granzyme B was still elevated in Treg-treated lung grafts on day 3 post-transplant compared to controls (supplementary figure 4n), but not in the dLNs (supplementary figure 4o).

At day 7 post-transplant, acute lung injury and lung allograft rejection scores (ISHLT A and B grades) were similar between animals receiving Treg-treated lungs and controls (supplementary figure 5a-b). CD4^+^ T cell numbers were reduced in Treg-treated lung allografts, although this observation did not reach statistical significance (figure 7a). Due to loss of tracking dye with time, our ability to detect transferred cells at day 7 was impaired. Nevertheless, FoxP3 expression in CD4^+^ T cells remained higher in Treg-treated allografts, compared to controls at this point (figure 7b). CD90 is upregulated on activated rat T cells [18], and in keeping with this finding, we observed fewer activated CD90^+^ CD4^+^ T cells in the lung allograft at day 7 post-transplant (figure 7c); reductions in CD90^+^CD4^+^ T cells were also seen in the dLNs but these findings were not statistically significant. No difference in Treg-related transcripts was observed between Treg-treated lung grafts and controls at day 7 post-transplant (supplementary figure 5e).

**Figure 7.**
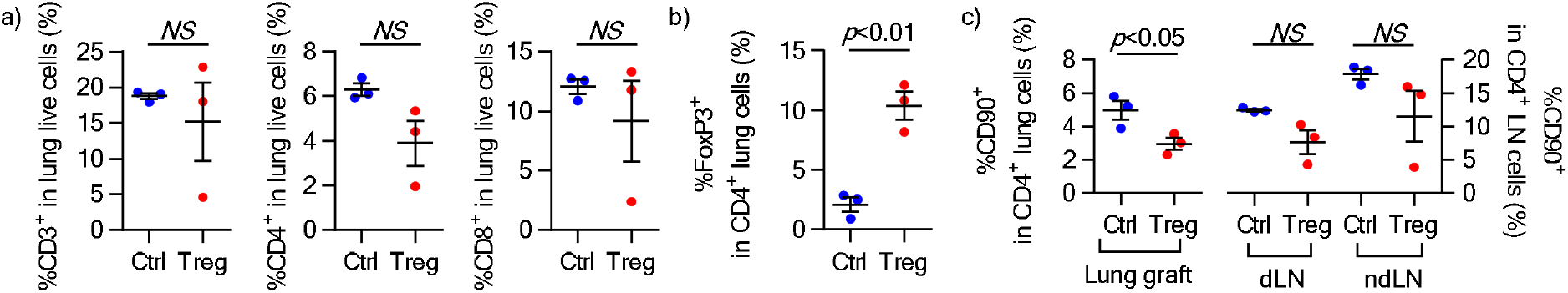
Control of T cell activation in the lung allograft at 7 days post-transplant. (a) Percentage of T cells in Treg-treated lung graft on day 7. Gating strategy is shown in supplementary figure 5c. (b) Percentage of FoxP3^+^ in CD4^+^ T cells was significantly increased in Treg-treated lung graft on day 7, compared to control lungs. (c) CD90 expression on CD4^+^ T cells was significantly decreased in the lung graft of Treg-treated animals compared to control cases, but not statistically significant in dLN and ndLN. Analysis of FoxP3 and CD90 expression in CD4^+^ T cells is shown in supplementary figure 5d. Treg, regulatory T cells; dLN, draining lymph node; ndLN, non-draining lymph node.

### Administration of allogeneic human Tregs during human EVLP

Human Tregs expanded 1,067.7 ± 363.4-fold and were 96.3 ± 1.2% viable after 21 days in culture (supplementary table 1). Viable cells were 81.4 ± 4.7% FoxP3^+^ and 81.8 ± 5.3% CTLA4^+^ (figure 8a). Characteristics of lung donors and EVLP parameters are shown in supplementary table 2. Injected Tregs were 87.1 ± 1.2% viable and 80.2 ± 6.7% CD4^+^CD127^-^. EVLP perfusate measurements are shown in supplementary table 1. Transferred Tregs were identifiable in perfusate and in lung tissue samples obtained 1h after injection (figure 8b-c). Expression of key functional Treg markers CTLA4, CD15s, CD39, 4-1BB, CCR4, and CXCR4 was higher in lung than in perfusate Tregs 1h after injection (figure 8d). However, there was no clear difference in FoxP3 or other molecules (figure 8e). In lung tissue, we observed an increase in Treg-related transcripts (IL-10, Granzyme B, and IDO-1) 1h after Treg injection compared to untreated control lungs over a similar time period on EVLP. Interestingly, FoxP3 and CTLA4 were not consistently upregulated in Treg-treated lungs compared to controls (supplementary figure 6d).

**Figure 8.**
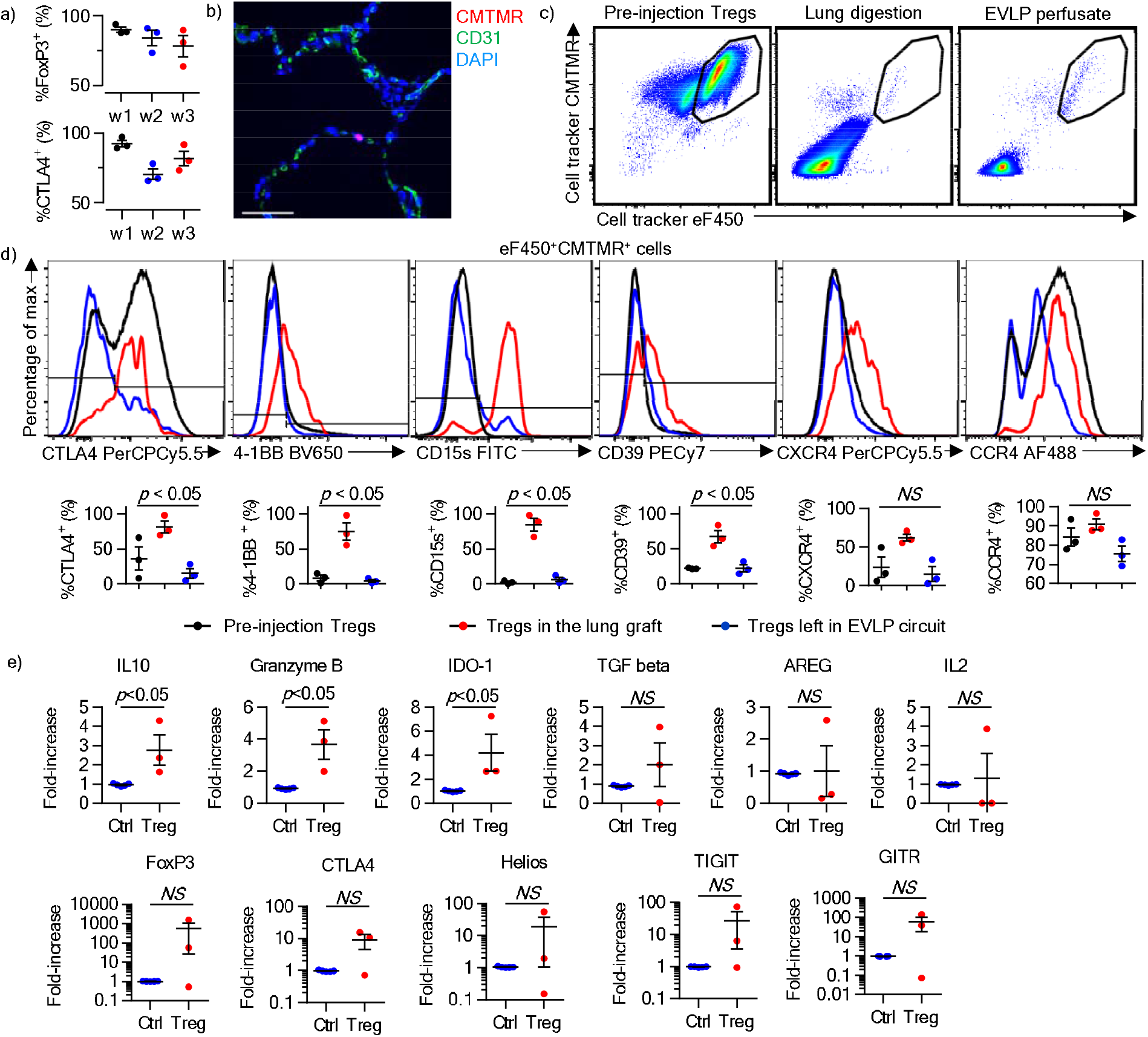
Delivery of human Tregs to allogeneic human lungs during EVLP. (a) Expression of FoxP3 and CTLA4 during 21 days of expansion. (b) Immunofluorescence staining of the lung graft 1 hour after Treg administration. CMTMR^+^ cells (red) were identifiable in the parenchyma. Images from each channel are shown in supplementary figure 6c. Scale bar = 50 µm. (c) Flow cytometric analysis of Tregs before injection (left panel), in the lung graft (middle panel) and in the perfusate (right panel) 1 hour after injection. (d) Analysis of the expression of the indicated markers on eF450^+^CMTMR^+^ cells in prior to injection (black lines), in the perfusate (blue lines) and in the lung allograft (red lines) 1 hour after injection, and those left in the EVLP circuit. There were significant differences in the expression of CTLA4, 4-1BB, CD15s, and CD39 on Tregs between the groups (repeated measures ANOVA). The expression of CXCR4 was relatively higher on Tregs in the lung graft than in those left in EVLP circuit (*p* = 0.0617, paired t test) although the intergroup difference was not significant. The majority of Tregs in the lung graft was positive for CCR4 (90.9 ± 2.8%). (e) Quantitative PCR analysis of the indicated transcripts in lung tissue. Transcripts measured at the end of EVLP and normalized to the housekeeping gene PPIA are displayed as a ratio to their abundance at the beginning of EVLP. Results from Treg-treated lungs (n = 3, red) are compared to results from untreated contemporaneous discarded lungs at the beginning and end of EVLP (blue lines). Mann-Whitney tests, *p* < 0.05 for IL-10, granzyme B and IDO-1. Treg, regulatory T cell; EVLP, *ex vivo* lung perfusion; PBMC, peripheral blood mononuclear cells.

## Discussion

Assessment of graft function prior to LTx using EVLP has expanded the donor pool, as it allows surgeons to transplant lungs that would previously have been considered unsuitable, with short- and long-term outcomes equivalent to conventional LTx [19]. Beyond these important advances, EVLP holds promise as a tool for delivery of advanced cell and gene therapies to reduce lung allograft injury [7, 20]. Furthermore, it provides the unique ability to immunologically manipulate the donor organ in the recipient ‘s favor, prior to the arrival of recipient leukocytes. With this concept in mind, we used EVLP to examine whether pre-transplant administration of expanded recipient Tregs to the graft could modify the alloimmune response at its earliest stages.

We were able to isolate and expand functionally suppressive rat CD4^+^CD25^high^ Tregs, similar to previous experiences with mouse and human Tregs [21, 22]. The cells did not cause injury or alter lung physiology across a wide range of doses and were taken up in a dose-dependent manner. About 25% of the administered cells remained in the EVLP circuit irrespective of the dose. Chemokine receptors including CCR4, CCR5 and CXCR3 can promote Treg entry into tissues in a variety of contexts [23]. We found that CCR4 and CXCR4, but not CCR5, CCD6, CCR7 or CXCR3, were more highly expressed on Tregs entering human lungs than those remaining in the EVLP circuit, suggesting that these receptors may be involved in migration of the cells into the lung. Whereas T cell receptor (TCR)-MHC interaction mediates CD8^+^ T cell uptake by allografts [24], its role in CD4^+^ T cell uptake is less clear [25, 26] and in any case, only ∼10% of T cells are alloreactive [27], suggesting that Treg entry into the lung was not strictly TCR-dependent. Whether TCR-MHC interaction and/or chemokine-chemokine receptor interaction is required for Treg uptake by allogeneic lungs during EVLP, or whether other mechanisms are involved, is unclear and is a subject of ongoing research.

Tregs taken up by the lungs entered the parenchyma, by a process that did not cause injury. Movement of water out of pulmonary capillaries into the lung interstitium and alveolar spaces contributes to lung injury and primary graft dysfunction after LTx [28, 29]. Typically, the alveolar-capillary barrier is disrupted, which can be revealed by staining for ZO-1 [7]. Although lung water was reduced in Treg-treated lungs compared with controls, we found no difference in ZO- 1 staining at the end of EVLP. This observation suggests that the degree of injury in these lungs may have been too mild to detect meaningful differences using this method. In keeping with this assertion, the wet-to-dry weight ratio of control lungs was also near normal [30]. It is conceivable that Tregs may have altered one or more physiologic alveolar fluid clearance pathways [31]. Whether polyclonally-expanded Tregs have a capacity for repair of more damaged lungs is an important question that awaits further investigation.

There was an increase in intra-graft Treg-associated transcripts (FoxP3, CTLA4, and GITR) at the end of EVLP in the rat model, and Tregs sorted from treated lungs retained suppressive function. The mildly reduced suppressive capacity of cells sorted from the lung might reflect a reduction in viability or loss of functional molecules as a result of enzymatic digestion, or could conceivably indicate that Tregs entering the lung were intrinsically less functional. Nevertheless, the transferred cells continued to exert immune regulation *in vivo* post-transplant. Tregs in the graft on day 3 exhibited increased proximity to antigen presenting cells (APCs), which are the principal site of immune regulation by Tregs *in vivo* [14]. Recipient Tconv, in contrast, displayed reduced proximity to APCs in the presence of Tregs, with Treg-associated APCs showing the lowest proximity to Tconv. Further, Tconv in Treg-treated lungs and dLNs exhibited reduced activation marker expression compared to controls. These are important findings, as it has been shown that CD11c^+^ APCs in the graft are the initial site of T cell priming in LTx, and that this process has already occurred by day 3 post-transplant [4]. We administered polyclonal allogeneic Tregs to human lungs and made a number of similar observations – including upregulation of immunoregulatory transcripts within the lungs following Treg administration.

Another advantage of our approach is the ability to deliver large numbers of Tregs directly to the site of alloreactive T cell activation. For polyclonal Treg therapies, such as the ones used here, it has been estimated that a 1:1 or 1:2 Treg:Tconv ratio is required within the target organ to control the recipient ‘s alloimmune response [32]. To achieve this ratio with systemic Treg administration might be difficult, but our strategy suggests a means by which to attain this goal.

Allograft-directed Treg therapy has limitations. We used polyclonal Tregs, so it is likely that only a minority of these cells were reactive to donor antigens. Ultimately, since donor antigen-reactive Tregs are more potent than polyclonal Tregs [33], it would be desirable to combine our approach with chimeric antigen receptor Tregs [34, 35]. We did not administer immunosuppressive drugs, and since it is known that calcineurin inhibitors can limit Treg function *in vivo* [36], it is likely that standard LTx immunosuppression would reduce the effectiveness of this approach. Additional work will be required to understand the impact of pre-transplant Treg therapy on allograft rejection. We were unable to track administered cells beyond day 3, which may have resulted from dilution of labeling dyes and/or cell death or migration; nevertheless, Foxp3^+^ cells were more prevalent among CD4^+^ T cells in Treg-treated grafts compared with controls at 7 days post-transplant. Clearly, however, since Foxp3^+^ cells only accounted for ∼10% of intragraft CD4^+^ T cells at day 7, further optimization will be required in order to inhibit rejection. Selective immunosuppressive approaches to enhance desired T cell responses – such as those mediated by Tregs – are emerging [37, 38] and it would be attractive to combine one of them with pre-transplant Treg administration.

In summary, recipient-derived expanded Tregs can be administered to rat and human lungs during EVLP. The cells did not injure the grafts and inhibited Tconv responses *in vitro* and *in vivo*. Our findings therefore support the concept that immune modulation can begin in the allograft prior to transplantation, thus opening the door to personalized organ-directed immunoregulatory cell therapy.

## Author contributions

S.J. conceived and designed the project. S.J., E.M., A.T., M.C., T.M., M.L., and S.K. contributed to the interpretation of data and critically revised the manuscript. E.M. and A.O. developed the method of rat Treg injection. E.M. and A.T. performed rat EVLP and transplantation. E.M., K.M.B., M.U. and B.J. isolated and expanded Tregs and performed suppression assays and flow cytometry. D.H and T.M. performed histologic grading of ALI in the lung graft. E.M., K.F.B., D.V., and Z.G. performed immunohistochemistry and immunofluorescence staining and analysis. Z.G. performed gene expression analysis. E.M. and S.J. contributed to the statistical analysis.

## Acknowledgments

We thank Toronto General Hospital EVLP team for supporting the human EVLP Treg injection experiments, and The Cell Science Research Foundation (Japan), The Kyoto University Foundation (Japan), and the International Society for Heart and Lung Transplantation for supporting E.M. via their research fellowship programs.

## Supplementary Appendix Material

Supplementary Materials and Methods

Supplementary figure 1

Supplementary figure 2

Supplementary figure 3

Supplementary figure 4

Supplementary figure 4 extended

Supplementary figure 5

Supplementary figure 6

Supplementary table 1

Supplementary table 2

Supplementary table 3

Supplementary table 4

Supplementary movie 1

