## Supplemental text for "*Ex vivo* delivery of regulatory T cells for control of alloimmune priming in the donor lung"

**Supplementary Materials and Methods**

*Purification of rat Tregs and T conv cells*

WKy rat cervical, axillary, inguinal, mesenteric, and mediastinal lymph nodes were crushed through 70 µm cell strainers to yield single cell suspensions. Cells were subjected to osmotic shock in 2 mL of NH_4_Cl RBC lysis buffer for 4 min at room temperature. After washing, cells were incubated with anti-CD8 (Abcam) and anti-CD20 antibodies (Thermo Fisher Scientific) for 10 min at 4^o^C. The cells were washed and incubated with anti-mouse IgG microbeads (Miltenyi Biotec) for 10 min at 4^o^C. Magnetically labelled cells were removed using an LD column (Miltenyi Biotec). CD8- and CD20-depleted cells labelled with anti-CD4 FITC and anti-CD25 PE antibodies prior to FACS sorting on a BD Biosciences FACSAria II or FACSAria III. As depicted in supplementary figure 1a, CD4^+^CD25^high^ cells, which corresponded to 1.2~1.6% of total cells, were sorted as Tregs. CD4^+^CD25^low^ cells were also collected as Tconv for Treg suppression assays (see below). A total of 0.221~0.478 × 10^6^ of sorted CD4^+^CD25^high^ cells were cultured at 0.025 × 10^6^ cells/mL in X-VIVO15 serum-free hematopoietic cell media (Lonza) with 5% fetal bovine serum (FBS, Gibco), 50 µM 2-mercaptoethanol (Thermo Fisher Scientific), 100 units/mL penicillin (Gibco), and 100 ug/mL streptomycin (Gibco), in the presence of 2500 IU/mL rhIL-2 (Proleukin, Chiron Therapeutics) and anti-rat CD3 and anti-rat CD28 coated microbeads (Miltenyi Biotec) at 1:40 cell-to-bead ratio. Tregs were expanded in culture for 7 days and were replenished on day 3 with fresh cell culture media containing 2500 IU/mL rhIL-2, diluting the cells to no more than 0.4 × 10^6^ cells/mL. The polyclonally expanded cells were harvested on day 7 and prepared for injection as described below.

*Details of rat Treg injection during rat EVLP followed by LTx*

F344 rats were intubated under anesthesia with 2% Isoflurane (Fresenius Kabi) and ventilated with 50% oxygen. At spine position, after flushing with 20 mL cold Perfadex (XVIVO Perfusion), the heart and lung bloc was extracted and connected to the EVLP circuit (Isolated Perfused Lung Systems, Harvard Apparatus, supplementary figure 1d-e) which was filled with 150 mL perfusate solution (Steen®, XVIVO Perfusion, Denver, CO) containing 10000 units Heparin, 50 mg methylprednisolone sodium, and 50 mg cefazolin sodium. Perfusion and ventilation commenced at 37^o^C under the following conditions: inspiratory/expiratory pressure: 9/4 cmH_2_O; I/E: 50%; respiratory rate: 40 per min; 21% oxygen. The perfusate was deoxygenated in a fiber membrane (Harvard Aparatus TYPE D150) filled with a gas mixture of 92% nitrogen and 8% CO2. The left atrial line pressure was maintained at 2 cmH2O against the hilum. At 60 min of EVLP, Tregs labelled with CellTracker Orange CMTMR Dye and/or eF450 cell tracker dye in 1 mL Steen or 1mL Steen containing no cells were injected into the EVLP circuit upstream of the lungs via the injection port. Perfusate was sampled just before Treg injection and at 5, 15, 30, 60, 120 and 180 min after Treg injection both upstream (PA port) and downstream (LA port) of the lung allograft. At 180 min after Treg injection, the heart and lung block was disconnected from the EVLP circuit and placed in cold Perfadex. After ventilating for less than 1 min with 50% oxygen and 2 cmH2O positive end-expiratory pressure (PEEP), the trachea was clamped. The left and right lungs were separated, keeping the left lung inflated, and the right lung was used for gene, histology, cytokine, flow cytometric analyses. The left lung was prepared for transplantation with cuff technique on cold Perfadex in 60 min after end of EVLP. Recipient WKy rats were anesthetized with 2% Isoflurane and ventilated with 21% oxygen and 2 cmH2O PEEP. After anastomosing the bronchus, pulmonary vein, and pulmonary artery, the lung graft was re-perfused 15 min after placing the graft into the left chest cavity, and the chest was closed. After recovering from anesthesia, the recipients were administered 10 ug/kg buprenorphine subcutaneously. During euthanasia at day 3 or day 7 post-transplant, recipients were anesthetized with 2% Isoflurane.

*In vitro rat Treg suppression assay*

Polyclonally expanded Tregs and freshly sorted Tconv were labelled with cell proliferation dye eFlour 450 (eF450, Invitrogen) and Vybrant CFDA SE Cell Tracer Kit (CFSE, Invitrogen) according to the manufacturer’s instructions, respectively. A total of 25 × 10^3^ of Tconv were co-cultured in duplicate with varying numbers of Tregs in AIM-V medium (Thermo Fisher Scientific) with 50 µM 2-mercaptoethanol, 100 units/mL penicillin, and 100 ug/mL streptomycin, in the presence of 1500 IU/mL rhIL-2 and anti-rat CD3/anti-rat CD28 coated microbeads at 1:2 of cell to bead ratio. After 96 hours, cells were stained for flow cytometric analysis on a BD Biosciences LSRII. The CD4^+^CFSE^+^ population in the eF450^-^FoxP3^-^ live cell gate was analyzed as Tconv as shown in supplementary figure 1c. Results were reported as percentage suppression, as described in figure 2b.

*Isolation and expansion of human Tregs*

Approximately 100 mL of whole blood was collected from a healthy donor, and PBMCs were isolated using Lymphoprep (STEMCELL TECHNOLOGIES) according to the manufacturer’s instructions. First, CD4^+^ cells were isolated by magnetic depletion of CD8^+^, CD14^+^, CD15^+^, CD16^+^, CD19^+^, CD36^+^, CD56^+^, CD123^+^, TCR^+^and CD235^+^ using a CD4^+^CD25^+^ T Cell Isolation Kit (Miltenyi Biotech). The isolated cells were labelled with anti-CD4, anti-CD25, and anti-CD127 antibodies and then CD4^+^CD25^+^CD127^low^ cells were isolated as human Tregs using FACS sorting on a BD FACSAria II or FACSAria III (supplementary figure 6a). Sorted Tregs were cultured in X-VIVO15 with 5% human AB serum in the presence of anti-human CD3/anti-human CD28 microbeads (Miltenyi Biotec) at a 1:2 cell-to-bead ratio and 500 IU/mL rhIL-2. The cells were expanded for 3 weeks during which they were replenished with 500 IU/mL rhIL-2 every 3~4 days diluting the cells to a density of 3-5 × 10^5^/mL. The cells were restimulated with anti-human CD3/anti-human CD28 microbeads (1:2 cell-to-bead ratio) at day 14. At day 21, when the cells had expanded ~1000-fold (1~2 × 10^9^ cells), Tregs were resuspended in 50 mL of freezing media (90% FBS and 10% dimethylsulfoxide) and stored in liquid nitrogen for 16-100 days until needed.

*Details of expanded human Treg injection during EVLP*

The lung was perfused with Steen solution at 37^o^C , and ventilated with 21% oxygen, 5 cmH2O PEEP, 8 mL/kg tidal volume (supplementary figure 6b), as previously described.^1^ After 3 hours of lung assessment on EVLP, at which time the decision not to transplant the lung was made by the clinical team on the basis of organ quality,^2^ cryopreserved Tregs were thawed, assessed for viability, and labelled with CMTMR and eF450 dyes according to the manufacturer’s instructions.

*Flow cytometry*

Rat lymphocytes were obtained by crushing spleen and lymph nodes through 70 µm cell strainers. Human and rat lung cells were prepared by digesting the lungs in 4.5 mL HBSS with 2% FBS, 10 mg/mL Collagenase A (Sigma Aldrich), and 10 mg/mL DNAse (Sigma Aldrich) with gentleMACS dissociator (Miltenyi Biotec) and subsequently filtering through 70 µm cell strainer. Clones and species for antibodies used for flow cytometry were described in Table S3. All samples were fixed using a transcription factor fixation and permeabilization buffer set (eBioscience) according to the manufacturer’s instructions and run on an LSRII. Data were analyzed with FlowJo software (FlowJo, LLC).

*Tissue section staining and histological assessment*

Tissue samples were fixed in 10% neutral buffered formalin and embedded in paraffin. 5 µm sections were stained with hematoxylin-eosin, immunohistochemical, or immunofluorescence stains. Clones and species for antibodies used for immunohistochemistry and immunofluorescence stains were described in Table S4. ImmPRESS™ Anti-rabbit IgG reagent (Vector Labs) and liquid DAB+ substrate chromogen solution (DAKO) were used for immunohistochemistry. Analysis of stained sections were performed by blinded observers (D.H. for ALI score, K.F.B. and S.J. for ZO-1 score, and D.V. for Tconv-Treg-APC analysis). ALI score (0-12) was assessed as a sum of 4 categories: alveolar space hemorrhage (0-3), vascular congestion (0-3), edema/fibrin (0-3), and interstitial white cells (0-3). Infiltration of the inflammatory cells to the peri-vascular area (A grade) and peri-bronchiole area (B grade) of the lung graft was assessed based on the criteria proposed by International Society of Heart and Lung Transplantation.^3^ For zonula occludens-1 (ZO-1) score, ZO-1 alveolar cell membrane staining intensity in 5 randomly selected images from each lung was determined in a semiquantitative manner (0-4), as previously described.^4^ The results were evaluated by two independent blinded investigators (K.F.B. and S.J.) and averaged. For Tconv-Treg-APC analysis, scanned images from 10 randomly selected locations from each lung were obtained. The average number of Tconv cells (CMTMR^-^CD3^+^) adjacent to MHC class II^+^ cells associated with, and not associated with, transferred Tregs (CMTMR^+^CD3^+^) was determined.

*Gene transcript analysis*

Lung and lymph node samples were put in RNAlater stabilization solution (Invitrogen) for 4~7 days and then stored in -80 ^o^C. Total RNA was extracted from the tissue using Tneasy Mini Kit (Qiagen) and converted into cDNA using an iScript Select Kit (Bio-Rad Laboratories, Hercules CA). SsoAdvanced Universal SYBR Green Supermix (Bio-Rad) and a Bio-Rad CFX384 Tough real-time PCR thermal cycler were used for PCR. Primers were designed across introns. Threshold cycle (Ct) values were determined using instrument software. Changes in gene expression were calculated using the 2^-ΔΔCt^ method with normalization to expression of peptidylprolyl isomerase A.

**Supplementary figure legends**

**Supplementary figure 1.** Rat Treg and EVLP procedures

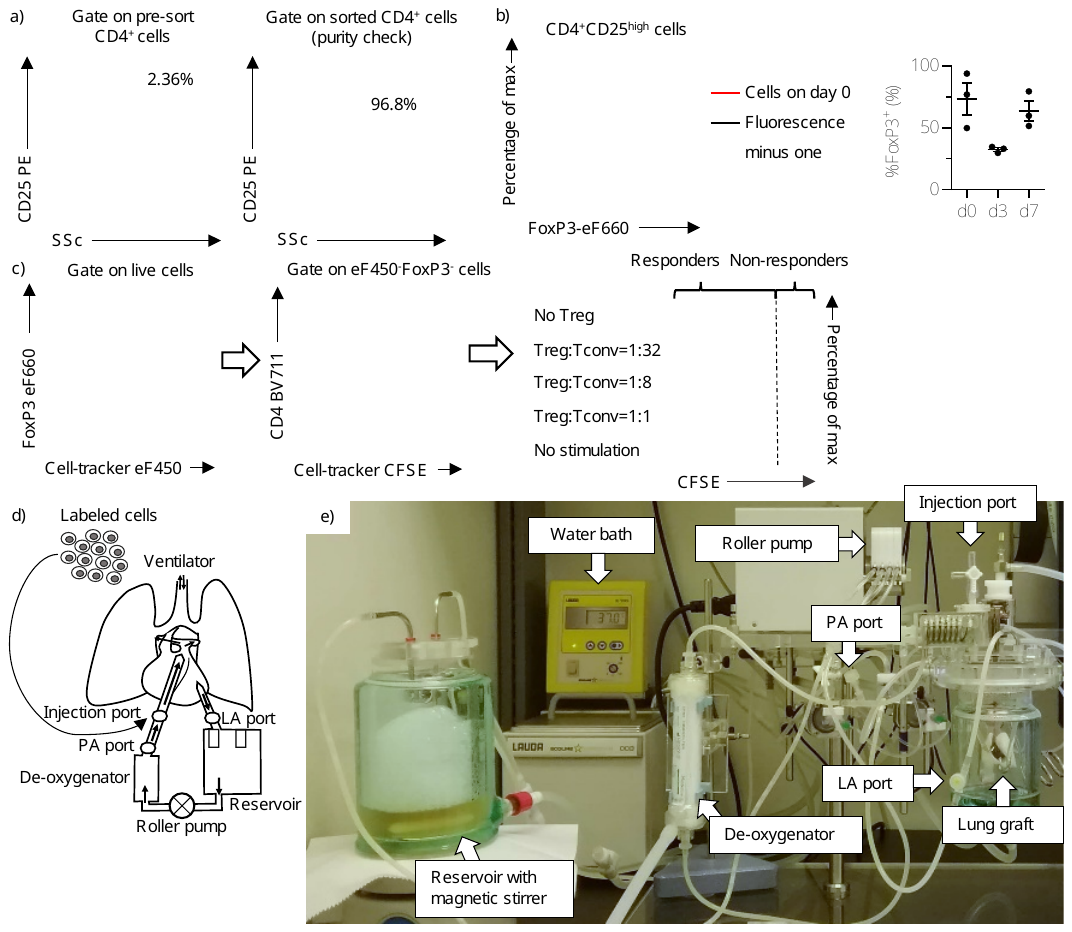

(a) WKy rat Treg isolation. After magnetic depletion of CD8^+^ and CD20^+^ cells, the top 2% of CD25^+^ cells in the CD4^+^ gate were sorted by FACS. 91.6 ± 1.4% of the sorted cells were positive for both CD4 and CD25 (n = 5). (b) 76.8% of the sorted cells were positive for FoxP3 (left panel). FoxP3 expression decreased on day 3 and then returned to the similar level of day 0 by day 7 (right panel). (c) Representative gating to identify conventional T cells (Tconv cells) at condition of Treg:Tconv = 1:32. eF450^-^FoxP3^-^CFSE^+^CD4^+^ population was considered as Tconv cells. In the right panel, histograms of CFSE intensity in WKy FoxP3^-^ CD4^+^ T cells (Tconv cells) following 4 days of culture with or without autologous expanded Tregs. Percent suppression was calculated as 100 × {(%Tconv with Treg − %Tconv without Treg)/%Tconv without stimulation}. (d) Expanded Tregs were injected in 1 mL of Steen solution at the injection port located upstream of the donor lungs. In control cases, 1 mL of Steen solution (no cells) was injected. Perfusate samples were collected from the PA and LA ports. After disconnecting the donor lungs from the circuit at the end of EVLP, the remaining perfusate in the circuit was collected. (e) Picture of rat EVLP system. See supplementary movie 1. Treg, regulatory T cell; EVLP, *ex vivo* lung perfusion; PA, pulmonary artery; LA, left atrium.

**Supplementary figure 2.** The effect of Treg injection during EVLP on the lung graft.

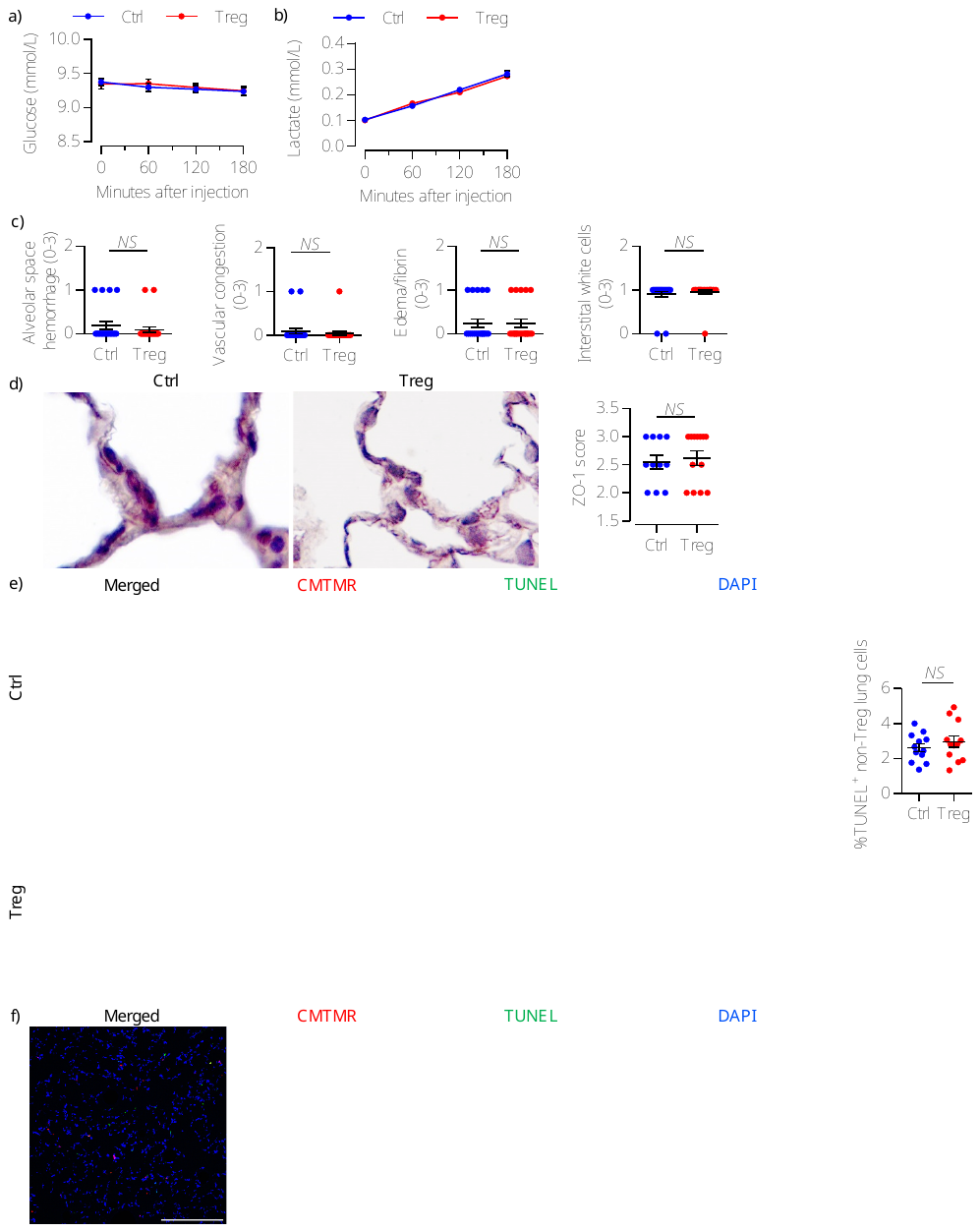

(a) Glucose level in the LA perfusate during EVLP. (b) Lactate level in the LA perfusate during EVLP. (c) Acute lung injury score (0-12) was an aggregate score which was derived by semi-quantification of alveolar space hemorrhage, vascular congestion, interstitial edema/fibrin deposition, and interstitial infiltration of white blood cells. (d) Representative pictures of ZO-1 staining (×100, *p*=NS). (e) TUNEL-positive apoptotic lung cells (green) in the lung after EVLP (*p*=NS, scale bars = 25 µm). (f) The percentage of TUNEL^+^ in CMTMR^+^ cells in the lung graft after EVLP was 0.97 ± 0.52% (n = 11, scale bar = 100 µm). CMTMR^+^ Tregs were not in the proximity of apoptotic cells. Treg, regulatory T cells; EVLP, *ex vivo* lung perfusion. LA, left atrium.

**Supplementary figure 3.** Kinetics of the injected cells in the recipient after lung transplantation.

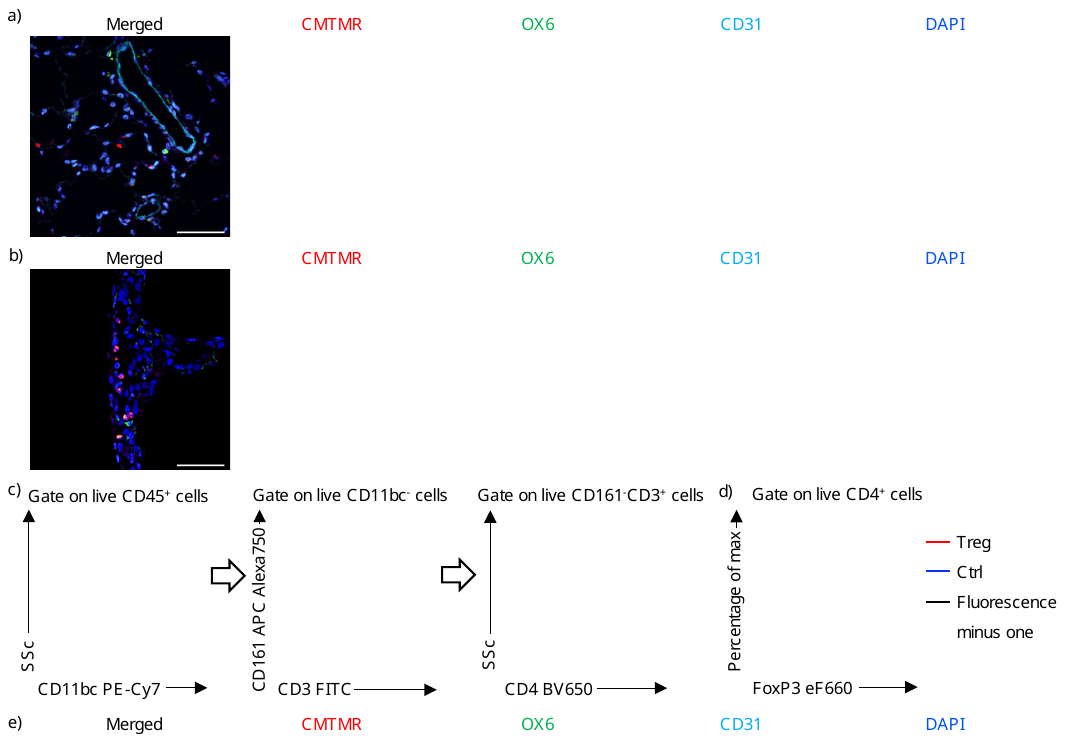

(a) Images show (left to right): merged image; CMTMR^+^ cells (red); OX6^+^ cells (rat MHC class II^+^ cells, green); CD31^+^ vessels (light blue); and nuclei (blue). (b) Images show (left to right): merged image; CMTMR^+^ cells (red); OX6^+^ cells (rat MHC class II^+^ cells, green); CD31^+^ vessels (light blue); and nuclei (blue). (c) Live CD45^+^CD11bc^-^CD161^-^CD3^+^CD4^+^ cells were considered as CD4^+^ T cells in flow cytometric analyses. (d) Analysis of FoxP3 expression in CD4^+^ in the lung graft on day 3. (e) Each channel visualized CMTMR^+^ cells (red), OX6^+^ cells (rat MHC class II^+^ cells; green), CD31^+^ vessels (light blue), or nuclei (blue). Treg, regulatory T cells; EVLP, *ex vivo* lung perfusion; MHC, major histocompatibility complex. Scale bars = 50 µm.

**Supplementary figure 4.** The immunomodulatory effect of Treg injection on the lung graft at day 3 post-transplant.

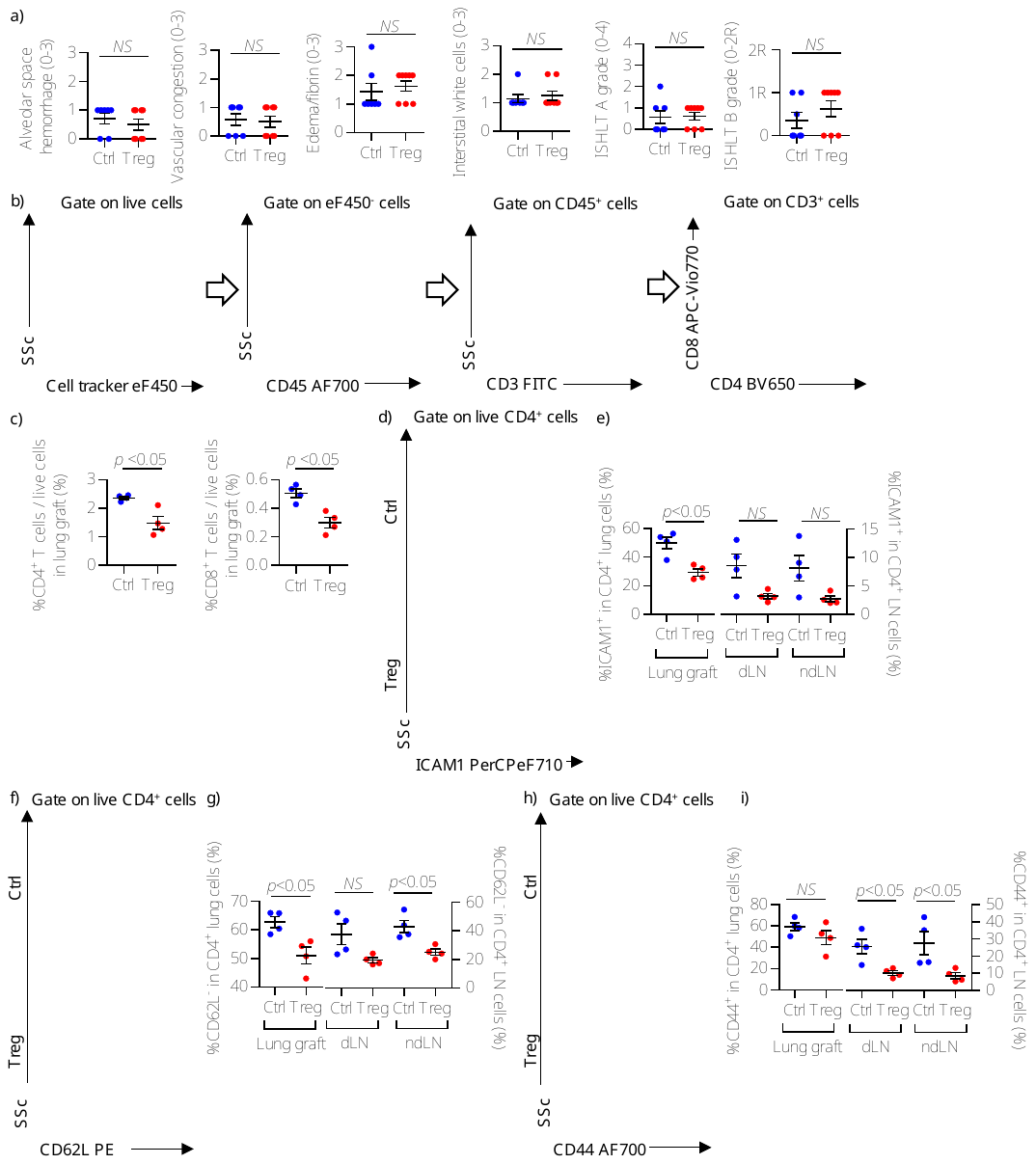

(a) Semi-quantification of alveolar hemorrhage, vascular congestion, interstitial edema/fibrin deposition, interstitial infiltration of white blood cells, ISHLT A grade (peri-vascular infiltration of inflammatory cells), ISHLT B grade (peri-bronchiole infiltration of inflammatory cells), and the percentage of inflammation area on day 3 post-transplant in Treg-treated and untreated lungs. (b) eF450^-^CD45^+^CD3^+^ cells were defined as non-transferred T cells. (c) The percentage of CD4^+^ (left panel) and CD8^+^ (right panel) cells in intra-graft live cells on day 3 was lower in Treg-treated animals than in controls (*p* < 0.05 and *p* < 0.05, respectively; Mann-Whitney U test). (d) Representative FACS dot plot to identify the ICAM1^+^ population in CD4^+^ T cells. (e) ICAM1 expression on CD4^+^ T cells was significantly decreased in lung grafts of Treg-treated animals compared to control cases, but not in lymph nodes. (f) Representative FACS dot plot to identify the CD62L^-^ population in CD4^+^ T cells. (g) The CD62L^-^ population in CD4^+^ T cells was decreased in the lung graft of Treg-treated animals compared to control cases, but not in their draining lymph nodes. (h) Representative FACS dot plot to identify CD44^+^ population in CD4^+^ T cells. (i) CD44 expression on CD4^+^ T cells was decreased in lymph nodes of Treg-treated animals compared to control cases, but not in the lung graft. EVLP, *ex vivo* lung perfusion; Treg, regulatory T cells; ISHLT, international society of heart and lung transplantation; dLN, draining lymph node; ndLN, non-draining lymph node.

**Supplementary figure 4 extended.**

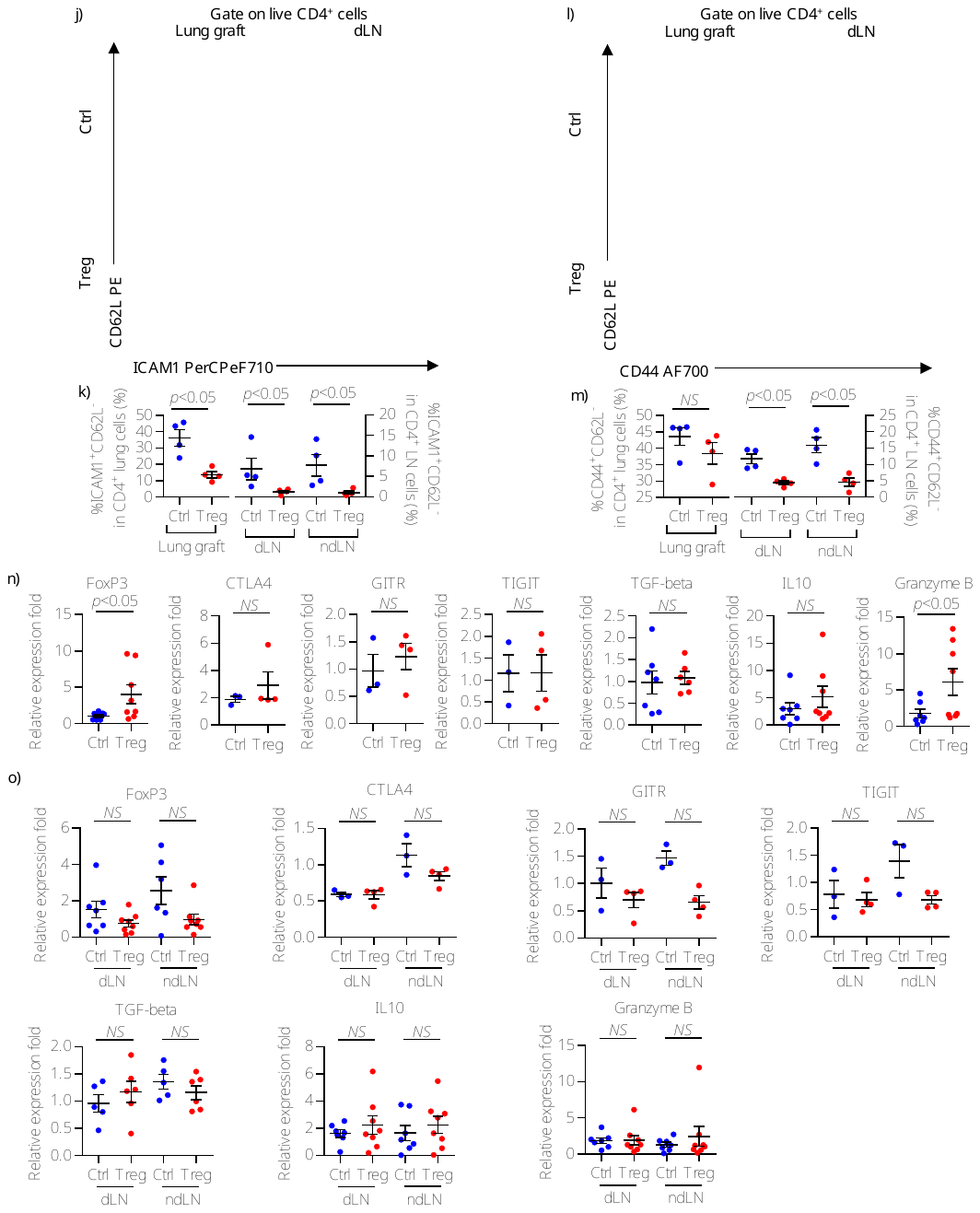

(j) Representative FACS dot plot to identify the ICAM1^+^CD62L^-^ population in CD4^+^ T cells. (k) The ICAM1^+^CD62L^-^ population of CD4^+^ T cells was decreased in the lung graft and lymph nodes of Treg-treated animals compared to control cases. (l) Representative FACS dot plot to identify the CD44^+^CD62L^-^ population in CD4^+^ T cells. (m) The CD44^+^CD62L^-^ population in CD4^+^ T cells was decreased in the lymph nodes of Treg-treated animals compared to control cases, but not in the lung graft. (n) Treg-related transcripts in the lung graft on day 3 post-transplant, represented as relative fold expression compared to the housekeeping gene PPIA. (o) FoxP3 expression in the lymph nodes of Treg-treated animals compared to control cases. Treg, regulatory T cells; EVLP, *ex vivo* lung perfusion; dLN, draining lymph node; ndLN, non-draining lymph node.

**Supplementary figure 5.** Administration of expanded Tregs during EVLP does not ameliorate lung allograft rejection at day 7 post-transplant.

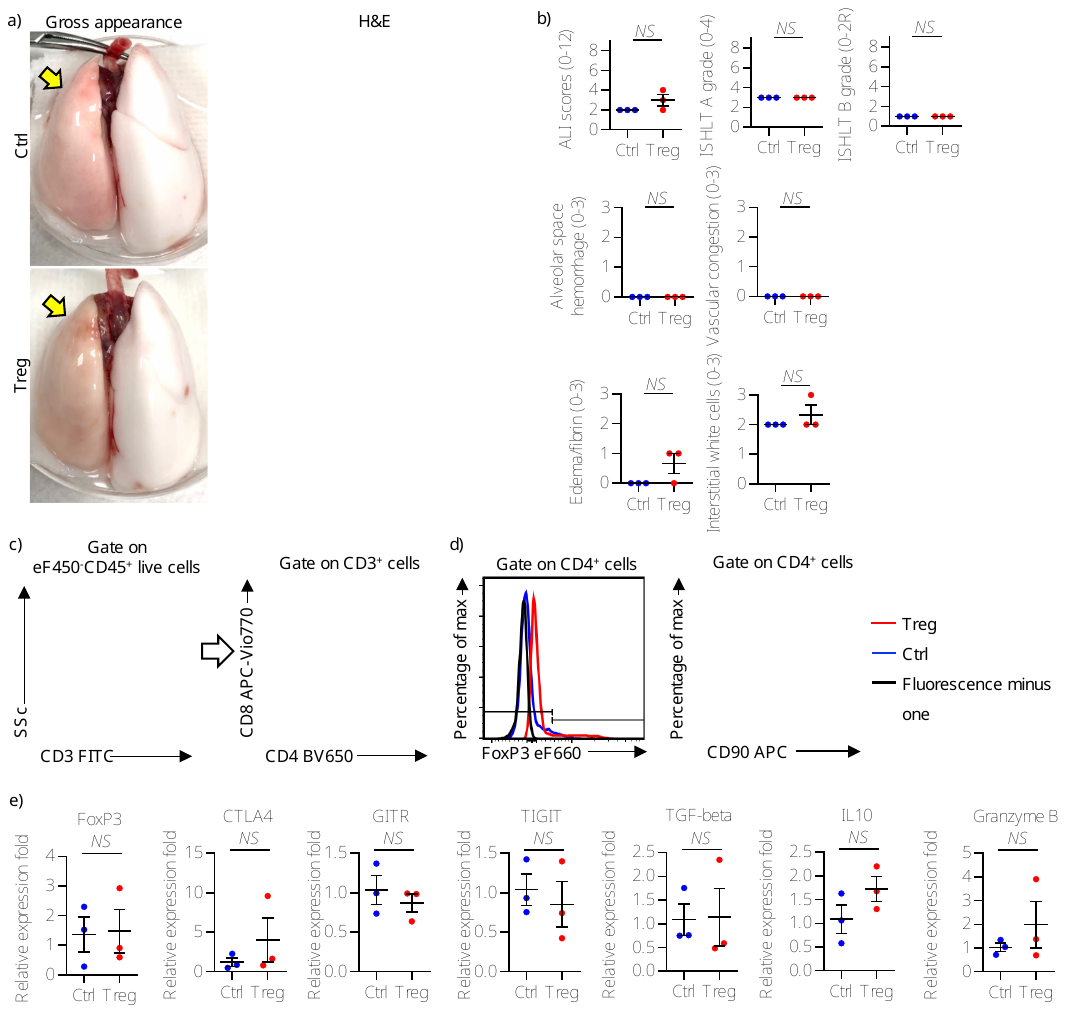

(a) Representative pictures of the gross appearance and the histology (hematoxylin and eosin staining; × 40) of the lung graft (yellow arrows) on day 7 post-transplant. (b) There were no intergroup difference in semi-quantification of acute lung injury (ALI) score which is the sum of alveolar space hemorrhage, vascular congestion, interstitial edema/fibrin deposition, and interstitial infiltration of white blood cells scores, ISHLT A grade, and ISHLT B grade in the lung grafts on day 7 post-transplant. (c) The representative FACS dot plot to identify CD4^+^ and CD8^+^ T cell populations in the lung graft. (d) Analysis of FoxP3 and CD90 expression in CD4^+^ T cells in the lung graft on day 7 post-transplant. (e) Treg-related transcripts in the lung graft on day 7 post-transplant, represented as a relative expression fold based on house-keeping gene. Treg, regulatory T cells; EVLP, ex vivo lung perfusion; ISHLT, International Society of Heart and Lung Transplantation; FACS, fluorescence activated cell sorting.

**Supplementary figure 6.** Delivery of human Tregs to allogeneic human lungs during EVLP.

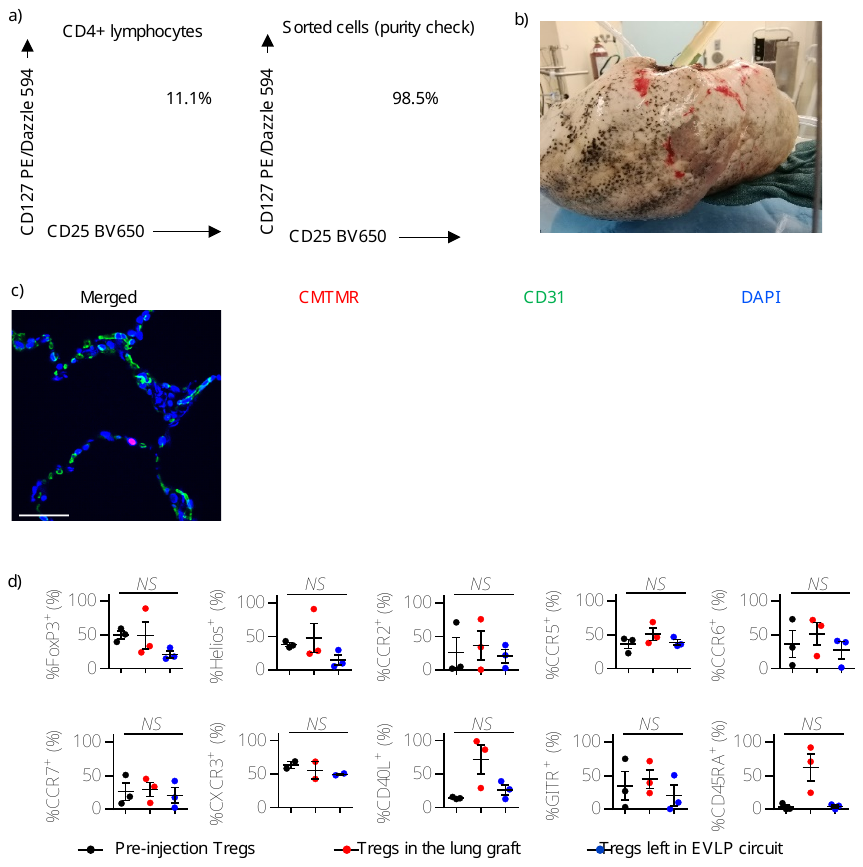

(a) CD4^+^CD25^high^CD127^low^ cells were sorted from CD4-enriched PBMC as human Tregs. Representative purity of the sorted cells is shown; overall 94.1 ± 2.2% of the sorted cells were found in CD25^+^CD127^-^ gate (n = 3). (b) Gross appearance of the donor right lung on EVLP. (c) Each channel visualized CMTMR^+^ cells (red), CD31^+^ vessels (green), or nuclei (blue). There was strong signal from DAPI channel since the administered Tregs were labelled with eF450 cell-tracker dye for flow cytometric analysis as well as CMTMR for IF staining. CMTMR^+^ Tregs were resident in the outside of CD31^+^ capillaries. Tregs, regulatory T cells. Scale bar = 50 µm. (d) Expression of FoxP3, Helios, CD40L, GITR, CD45RA, and chemokine receptors on Tregs taken from the lung and those remaining in the EVLP circuit 1h after Treg injection. Tregs, regulatory T cells; FACS, fluorescence activated cell sorting; EVLP, *ex vivo* lung perfusion; PBMC, peripheral blood mononuclear cells.

**Supplementary table 1.** The conditions of human Tregs and human lung EVLP

| Variables | EX1 | EX2 | EX3 |
| --- | --- | --- | --- |
| Number of sorted Tregs on day 0 | 1.40 × 10^6^ | 1.30 × 10^6^ | 1.87 × 10^6^ |
| Viability on day 21 before cryopreservation (%) | 98.0 | 94.0 | 97.0 |
| Duration of cryopreservation (days) | 80 | 121 | 38 |
| Number of Tregs injected (n) | 0.5 × 10^9^ | 0.8 × 10^9^ | 0.4 × 10^9^ |
| Viability at injection (%) | 87.2 | 89.0 | 85.8 |
| Fold-expansion in 21 days | 1307.1 | 1538.5 | 352.9 |
| Number of Tregs injected (n) | 0.5 × 10^9^ | 0.8 × 10^9^ | 0.4 × 10^9^ |
| Viability at injection (%) | 87.2 | 89.0 | 85.8 |
| PaO2/FiO2 in EVLP perfusate just before Treg injection (mmHg) | 406.4 | 373.0 | 433.7 |
| PaO2/FiO2 in perfusate 1h after Treg injection (mmHg) | 406.1 | 346.8 | 339.3 |
| Treg, regulatory T cell; PaO2/FiO2, ratio of the partial pressure of oxygen in the perfusate (PaO2) to the fraction of inspired oxygen (FiO2). | | | |

**Supplementary table 2**. Clinical features of donor lungs in the human EVLP experiment.

| Variables | | Treg injection (n = 3) | Ctrl (n = 5) |
| --- | --- | --- | --- |
| Lung donor sex (female/male, n) | | 0/3 | 2/3 |
| DCD/DBD (n) | | 2/1 | 3/2 |
| EVLP type (n) | | 2/1 | 3/2 |
|  | Rt. Single lung EVLP | 2 | 2 |
|  | Lt. single lung EVLP | 1 | 0 |
|  | Double lung EVLP | 0 | 3 |
| Treg, regulatory T cell; Ctrl, control lungs for OCR analysis; DCD, donation after cardiac death; DBD, donation after brain death. | | | |

**Supplementary table 3.** Antibodies for flow cytometry

| Marker | Fluorochrome | Reactivity | Company | Clone | Dilution |
| --- | --- | --- | --- | --- | --- |
| CD3 | FITC | Rat | BD Biosciences | G4.18 | 1/100 |
| CD4 | FITC | Rat | eBioscience | OX35 | 1/100 |
| CD4 | BV650 | Rat | BD Biosciences | OX-35 | 1/100 |
| CD4 | BV711 | Rat | BD Biosciences | OX-35 | 1/100 |
| CD25 | PE | Rat | eBioscience | OX39 | 1/100 |
| CD25 | BV650 | Rat | BD Biosciences | OX-39 | 1/100 |
| CD8a | PerCP-eF710 | Rat | eBioscience | OX8 | 1/100 |
| CD8a | APC-Vio770 | Rat | Miltenyi Biotec | REA437 | 1/100 |
| CD11b/c | PE-Cy7 | Rat | BD Biosciences | OX-42 | 1/100 |
| CD44 | AF700 | Rat | Novus biologicals | OX-50 | 1/100 |
| CD45 | AF700 | Rat | Biolegend | OX-1 | 1/100 |
| CD45 | BV711 | Rat | BD Biosciences | OX-1 | 1/100 |
| CD62L | PE | Rat | BioLegend | OX-85 | 1/100 |
| CD90 | APC | Rat | BioLegend | OX-7 | 1/100 |
| CD161 | APC-Vio770 | Rat | Miltenyi Biotec | REA227 | 1/100 |
| CTLA4 | Biotin | Rat | Thermo/invitrogen | WKH 203 | 1/100 |
| ICAM1 | PerCP-eF710 | Rat | eBioscience | 1A29 | 1/100 |
| Rat Fc Block (anti-CD32) | - | Rat | BD Biosciences | D34-485 | 1/100 |
| FoxP3 | PE-eF610 | Rat and human | eBioscience | FJK-16s | 1/50 |
| FoxP3 | eF660 | Rat and human | eBioscience | 150D/E4 | 1/50 |
| Viability | Fixable viability stain 700 | Rat and human | BD Biosciences | - | 1/1000 |
| Viability | Fixable viability stain PE-CF594 | Rat and human | BD Biosciences | - | 1/1000 |
| CD4 | BV711 | Human | BD Biosciences | SK3 | 1/100 |
| CD8 | APC/Cy7 | Human | Biolegend | SK1 | 1/100 |
| CD25 | BV785 | Human | Biolegend | BC96 | 1/100 |
| CD127 | PerCP-Cy5.5 | Human | Biolegend | A019D5 | 1/100 |
| GITR | APC/Fire750 | Human | Biolegend | 108-17 | 1/100 |
| Helios | PE/Cy7 | Human | Biolegend | 22F6 | 1/50 |
| CD40L | Biotin | Human | Biolegend | 24-31 | 1/100 |
| CD45RA | FITC | Human | Biolegend | HI100 | 1/100 |
| CTLA4 | PerCP-Cy5.5 | Human | Biolegend | BNI3 | 1/100 |
| 4-1BB | BV650 | Human | Biolegend | 4B4-1 | 1/100 |
| CD15s | FITC | Human | Biolegend | FH6 | 1/100 |
| CD39 | PE/Cy7 | Human | Bioledgend | A1 | 1/100 |
| CCR2 | BV605 | Human | Biolegend | k036C2 | 1/100 |
| CCR4 | AF488 | Human | R&D Systems | 205410 | 1/100 |
| CCR5 | APC/Cy7 | Human | Biolegend | J418F1 | 1/100 |
| CCR6 | Biotin | Human | BD Biosciences | 11A9 | 1/100 |
| CCR7 | PE-Cy7 | Human | BD Biosciences | 3D12 | 1/100 |
| CXCR3 | BV 785 | Human | Biolegend | G025H7 | 1/100 |
| CXCR4 | PerCP-Cy5.5 | Human | Biolegend | 12G5 | 1/100 |
| Human Fc Block | - | Human | BD Biosciences | - | 1/100 |

**Supplementary table 4.** Antibodies for IF and IHC

| Marker | Host species | Reactivity | Company | Dilution | Application |
| --- | --- | --- | --- | --- | --- |
| OX-6 | Mouse | Rat | BD bioscience | 1/200 | IF |
| CD31 | Rabbit | Rat and human | abcam | 1/300 | IF |
| CD3 | Rabbit | Rat | DAKO | 1/200 | IF |
| ZO-1 | Rabbit | Rat | Invitrogen | 1/300 | IHC |
| IF, immunofluorescence staining; IHC, immunohistochemistry staining | | | | | |

**Supplementary movie 1.** Rat heart-lung block perfused and ventilated during EVLP.
